# Structure of an RNA polymerase ribozyme replication complex

**DOI:** 10.64898/2026.08.30.748161

**Authors:** Timothy S. Strutzenberg, David P. Horning, Wesley G. Cochrane, Leonardo Andrade, Xuan Han, Gerald F. Joyce, Dmitry Lyumkis

## Abstract

Life began with the emergence of a molecule that could replicate its own genetic material, a task plausibly mediated by an RNA-dependent RNA polymerase ribozyme. Here, we present the structure of such a polymerase ribozyme, bound to RNA substrates comprising the template, primer, and nucleoside triphosphate (NTP) analog. The structure reveals how directed evolution shaped flanking elements around a highly conserved catalytic core derived from the ancestral class I ligase ribozyme. Each element serves as a functional module, positioning the primer-template duplex and incoming NTP within the active site of the enzyme. This emergent domain organization is remarkably similar to the “right hand” configuration of polymerase proteins, suggesting a common functional form for copying nucleic acids, regardless of biopolymer catalyst.

## Main Text

The most fundamental process of life is the replication and evolution of genetic material. Replication is performed by polymerase proteins that copy a DNA or RNA template to generate complementary nucleic acid products, utilizing (deoxy)nucleoside 5′-triphosphates (dNTPs or NTPs) as building blocks. In the early history of life, prior to the origin of encoded protein synthesis, RNA enzymes (ribozymes) that catalyze the RNA-templated synthesis of RNA may have replicated primordial RNA genomes (*1, 2*). Although such ribozymes have not been shown to exist within contemporary biology, they have been generated by directed evolution starting from a pool of random-sequence RNAs and refined by selective amplification (*3–12*).

The *in vitro* evolved class I ligase ribozyme catalyzes the RNA-templated ligation of two RNA substrates in a reaction analogous to NTP addition to the 3′ end of an RNA primer (*3, 4*). A high-resolution crystal structure of the ligase demonstrated that it folds into a tripod-like structure, with three coaxially-stacked helical legs that converge near the site of ligation (*13, 14*). The catalytic site is composed of two backbone phosphates that position a catalytic Mg^2+^ ion, together with two co-stacked cytosines that assist in catalysis.

Employing directed evolution methods, the class I ligase was evolved to function as an RNA polymerase, initially with the ability to add up to 14 nucleotides to a template-bound RNA primer (*6*). Further evolution enabled the polymerase to add many dozens of nucleotides (*8*). Two major innovations led to this substantial improvement in activity. The first involved addition of a 76-nucleotide random-sequence region that evolved to form an “accessory domain” to assist in sequence-general recognition of the RNA template and NTP substrate (*6, 7*). The second was the emergence of Watson-Crick (WC) pairing between the 5′ end of the ribozyme and 3′ end of the template, termed the “processivity tag”, which ensures high effective concentration of the primer-template relative to the enzyme (*8*).

A construct that contains both the accessory domain and processivity tag, named tC19Z, was the first polymerase ribozyme that could synthesize other functional RNAs (*8*). It also was used to initiate an extended lineage of 85 successive generations of directed evolution (*9–12*), culminating in the 85h34 polymerase that is the subject of the present study. This directed evolution campaign sought to optimize the efficiency, sequence generality, and fidelity of polymerization by requiring the polymerase to synthesize ever larger and more complex functional products, including aptamers and ribozymes.

Directed evolution led to a >600-fold improvement in polymerase activity and a more general capacity to copy diverse RNA sequences. By the 52nd round of evolution, a major structural rearrangement had occurred in the polymerase, resulting in a novel tertiary element that lies in close proximity to the active site (*11*). Consequently, the polymerase no longer depended on the processivity tag and was able to engage the primer-template complex with micromolar affinity through tertiary interactions (*15*). During rounds 52 through 85, emphasis was placed on improving polymerase fidelity, employing reduced concentrations of Mg^2+^ and requiring the polymerase to synthesize a functional class I ligase (*12*). Consequently, the accuracy of polymerization for synthesis of the class I ligase increased from 87% per nucleotide at round 52 to 95% per nucleotide at round 85.

### The structure of the 85h34 polymerase holoenzyme

The improved properties of 85h34, including its enhanced stability at lower concentrations of Mg^2+^, provided an opportunity to capture a high-resolution structure of the holoenzyme, including the ribozyme, template, primer, and incoming NTP. A 42-nucleotide template was designed that binds a complementary 20-nucleotide primer, followed by a short downstream template region that includes the template portion of the processivity tag. In the presence of all four NTPs, 85h34 rapidly extends the primer to full length within 10 seconds (fig. S1A). Analysis by mass photometry showed that the ribozyme, template, and primer form a well-behaved trimolecular complex (fig. S1B). To capture the incoming NTP, a non-reactive α,β-methyleneadenosine 5′-triphosphate analog was provided, which binds to a template uridine immediately adjacent to the 3′ end of the primer. The *holo*-85h34 complex was subjected to cryo-EM single-particle analysis and a map was derived from 1.15 million particles, resolved globally to 2.8 Å (fig. S2; fig. S3). The corresponding atomic model revealed the structure of this complex (table S1).

Over the course of many generations of directed evolution, the ancestral class I ligase core of the polymerase was elaborated by the emergence of novel functional domains (Fig. 1). A new pseudoknot pairing, named P8, creates a perpendicular extension from the P3-P6-P7 leg of the ligase core to meet the processivity tag at the 5′-end of the polymerase. This helical extension crosses over the minor groove of the primer-template duplex, positioning conserved J1/3 nucleotides in a similar manner as in the ligase active site. At the other end of the ribozyme, the 3′-accessory domain (A3, AJ3/4, and A4) drapes over the catalytic core, positioning the AJ3/4 loop directly over the 3′ face of the incoming nucleotide within the active site.

**Fig. 1.**
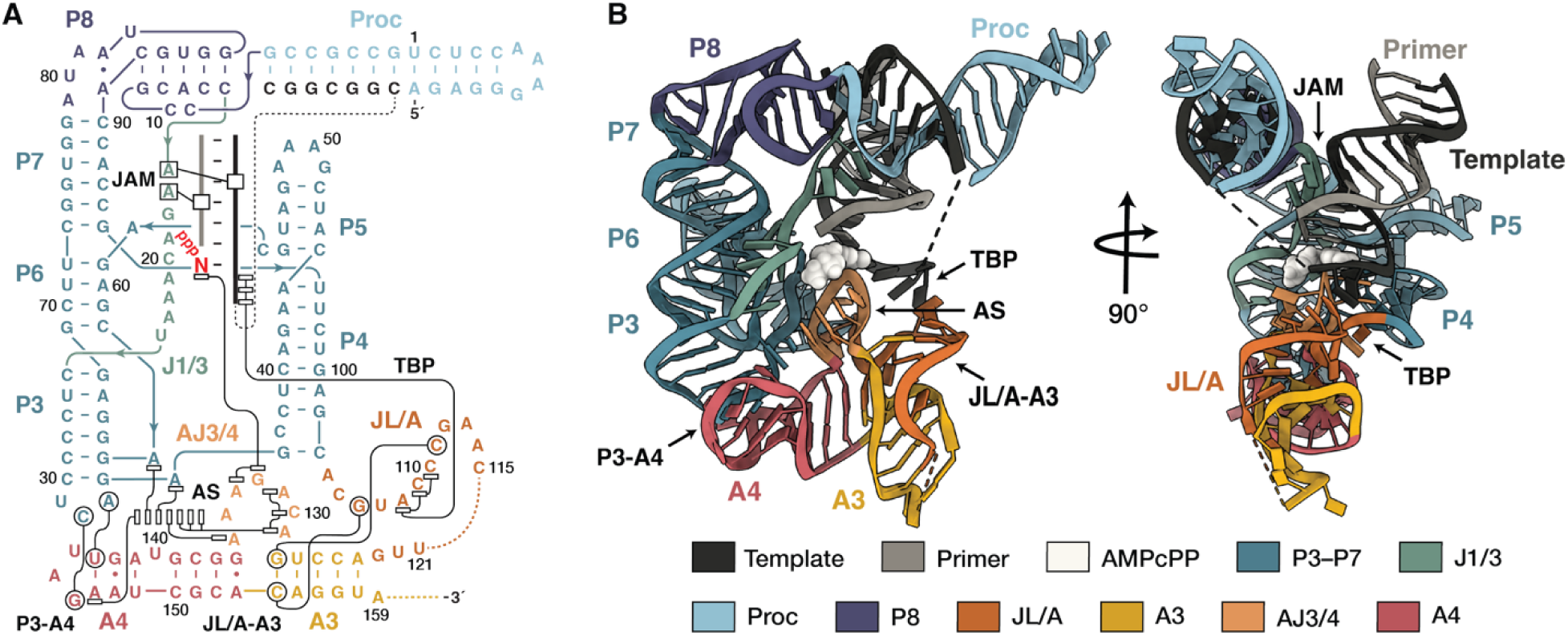
Structural basis of RNA polymerization by the 85h34 ribozyme. (**A**) Secondary structure diagram and (**B**) atomic model derived from a cryo-EM reconstruction of the 85h34 holoenzyme, colored according to the legend. The non-hydrolyzable α,β-methylene ATP analogue (AMPcPP) is shown as space-filling atoms at the center of the complex. Paired regions, including the processivity tag (Proc), and other points of interest are labeled. Notable tertiary contacts are shown as black lines in the secondary structure. These features include the J1/3 A-minor (JAM) interactions with the primer-template, the adenine stack (AS), the template base-stacking platform (TBP), and tertiary contacts between P3 and A4 and between JL/A and A3.

Four AJ3/4 nucleotides (A131, C130, A129, and A135) contribute to a prominent seven-nucleobase stack extending from A4 in the 3′-accessory domain to the active site. All but one of these nucleobases are adenines. We accordingly term this unique structural feature the “adenine stack”. The stack is formed by the four nucleotides from AJ3/4, two extra-helical adenylates from the ligase core (A35 and A63), and an adenylate from the A4 stem of the accessory domain (A147). A131 and C130, which are at the top of the adenine stack, lie parallel to the β,γ-pyrophosphate of the incoming NTP and are anchored to stems P3 and P4 of the core by tertiary interactions with A4 and A3, respectively. In addition, the JL/A connector, which links the ligase and accessory domains, interacts with the downstream single-stranded portion of the template through a base-stacking platform.

### A conserved catalytic center and modified NTP pocket

Most of the catalytic core (P3-P6-P7 helix, P4-P5 helix, and active site J1/3 nucleotides) has been conserved, deviating by less than 1 Å from the class I ligase and preserving two legs of the original tripod (Fig. 2A). The third leg (consisting of P1 and P2) has been replaced by the primer-template duplex, capped by the AJ3/4 and JL/A portions of the accessory domain to complete the active site. In the ligase structure, the P2 stem stacks over the 5′-terminal GTP, positioning it for ligation. In 85h34, the JL/A strand, which forms half of P2 in the ligase, is shifted by more than 4 Å to accommodate a new nucleotide binding pocket formed by AJ3/4. This configuration is consistent with prior crosslinking studies that placed these two regions near the template position opposite the incoming nucleotide (*15*).

**Fig 2.**
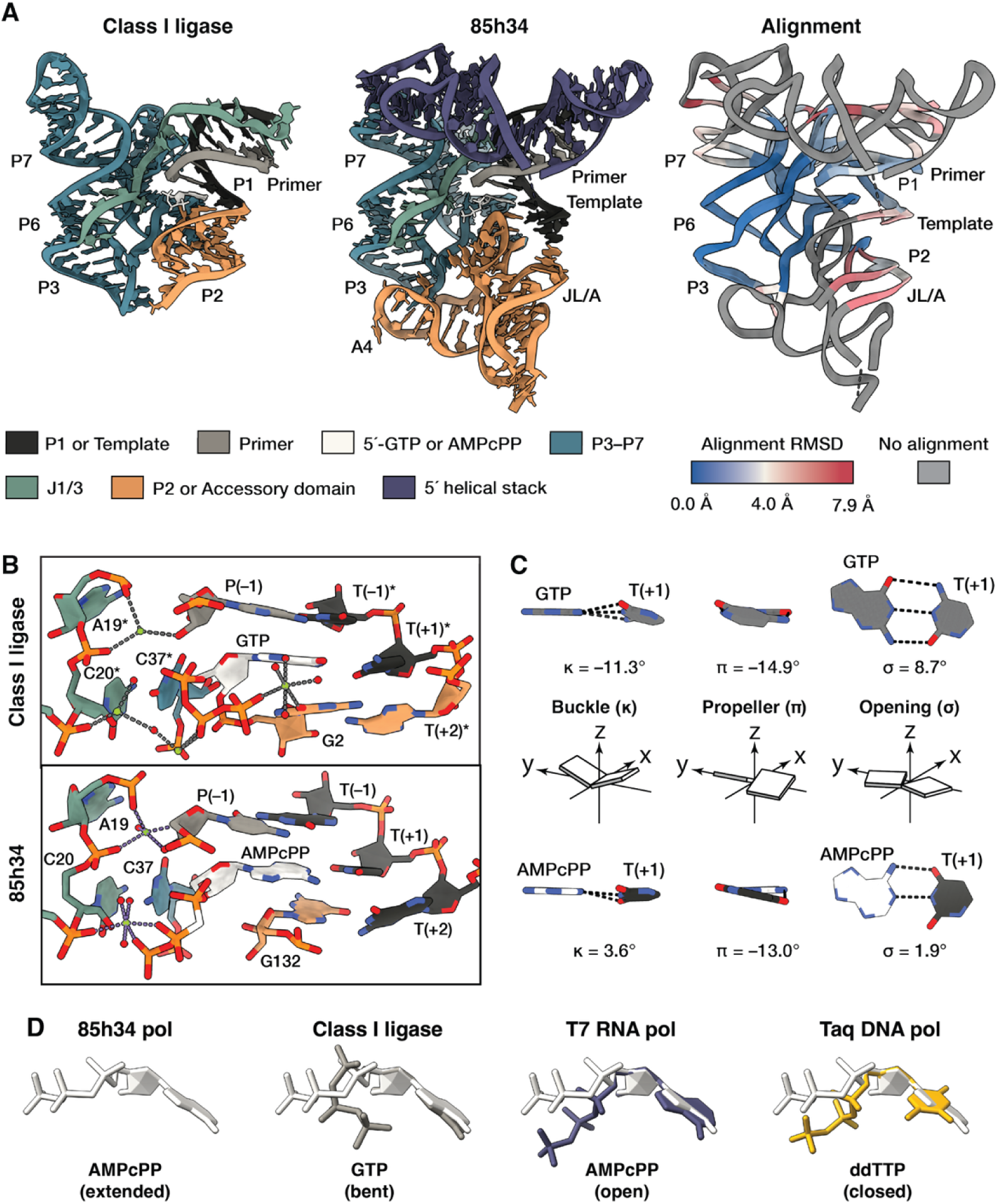
Evolution results in reconfiguration of the active site. (**A**) Atomic models comparing the class I ligase pre-ligation complex and 85h34 replication complex, colored according to the legend. The root mean squared deviations (RMSD) between the class I ligase and 85h34 are painted onto the ribbon depiction of 85h34 according to the color key. (**B**) Atomic coordinates of eight nucleotides around the incoming NTP. Asterisks indicate adaptation of the ligase numbering convention to match that of 85h34. Note that C37* was mutated to uridine to obtain the structure of the ligase (*13*). (**C**) Base-pairing geometry of the incoming NTP was calculated using the 3DNA webserver (*29*), with atomic coordinates shown for both the class I ligase and 85h34. (**D**) Conformation of the incoming NTP in the 85h34 ribozyme, class I ligase (PDB 3R1L) (*14*), T7 RNA polymerase (PDB 1S76) (*30*), and Taq DNA polymerase (PDB 1QTM) (*31*). NTPs were aligned with the ribose of AMPcPP in 85h34.

Around the active site, the key catalytic nucleotides are conserved, even as binding to the incoming nucleotide has changed. As first observed in the pre-ligation complex of the class I ligase (*13, 14*), the phosphates of nucleotides 19–21 coordinate Mg^2+^, nucleobases C20 and C37 stack over the ribose of the incoming NTP, and C37 provides additional hydrogen-bonding donors to the triphosphate (Fig. 2B). G132 from the AJ3/4 loop occupies a similar position as G2 in the ligase P2 stem, located over the substrate nucleobase, but in a distinct orientation, presenting ribose O4′ and the nucleobase to the incoming nucleotide and its complement. These interactions appear to improve co-planar WC-pairing geometry of the incoming nucleotide and template in 85h34 compared to the class I ligase (Fig. 2C). Increased stringency of WC pairing is a known fidelity mechanism, first described for protein polymerases (*16*). In the ligase, the triphosphate bends over the GTP ribose, with the γ-phosphate coordinated to a Mg^2+^-4H_2_O cluster bound to the Hoogsteen face of the 5′-terminal GTP and adjacent G2 nucleobase. The orientation of the triphosphate in 85h34 is more like that seen in polymerase proteins (Fig. 2D), extending away from the ribose, with the β,γ-phosphates coordinating a second Mg^2+^ that is bound to A21. Despite these observations, the precise mechanisms of catalysis and discrimination against mis-paired NTPs cannot be determined from this structure alone (for further discussion, see supplemental note).

### Organization of the NTP pocket and unpaired template nucleotides by the accessory domain

The accessory domain forms the NTP binding pocket by orienting the terminus of the adenine stack and adjacent AJ3/4 nucleotides over the incoming nucleotide. This is achieved by flanking tertiary anchors to the A3 and A4 stems of the accessory domain, each involving a “base-swap”, in which a paired nucleotide within a stem is replaced by another nucleotide (Fig. 3A). A base-swap with the A4 stem establishes the A147 nucleobase at the floor of the adenine stack. C32 and A33, which lie at the apex of the P3 stem, form WC and reverse Hoogsteen pairs to G146 and U143, respectively. Rather than the expected pairing to U143 within the A4 stem, A147 is ejected from the helix and stacks with A63 and A35 in the catalytic core, which in turn bridge P6 to P3 and P3 to P4, respectively.

**Fig. 3.**
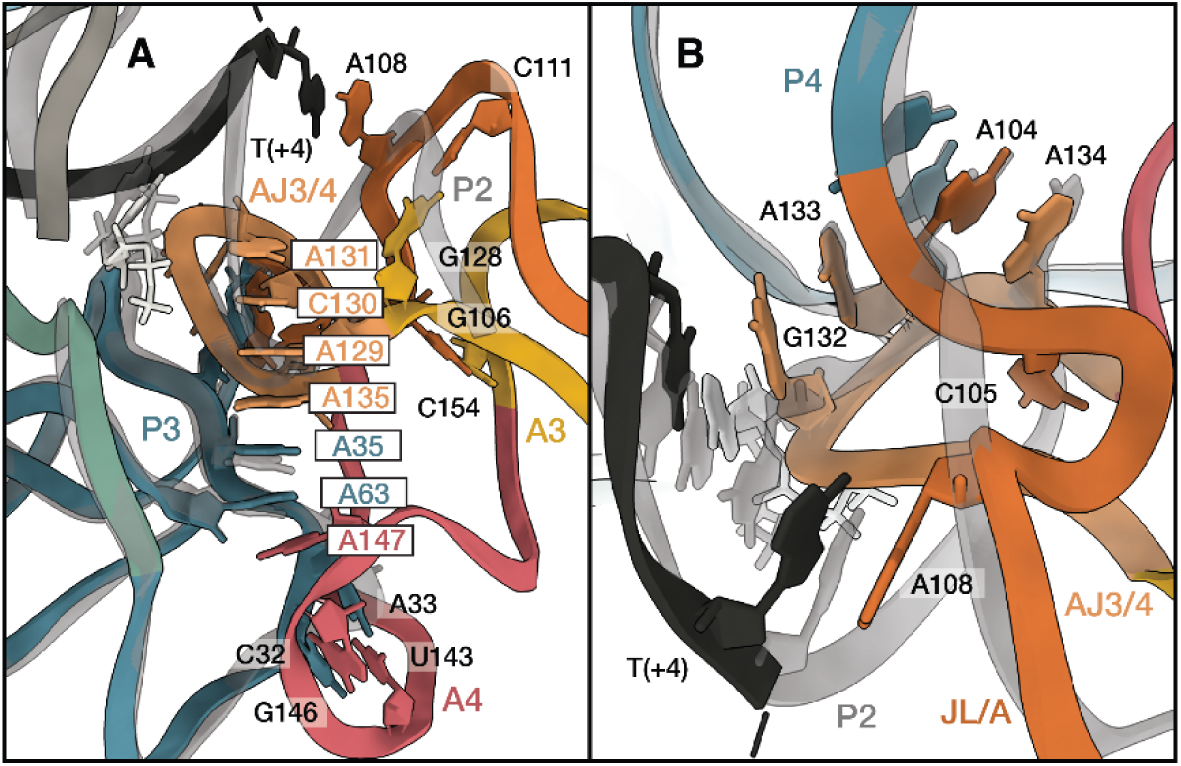
Tertiary contacts of the accessory domain. Ribbon diagrams of the 85h34 accessory domain (colored as in Fig. 1), aligned with class I ligase (colored gray). Notable residues are displayed and labeled. Asterisks indicate adaptation of the ligase numbering convention to match that of 85h34. Focused views are shown of (**A**) the adenine stack, and (**B**) AJ3/4 nucleotide interactions.

At the portion of the A3 stem that lies adjacent to the AJ3/4 loop, G106 of the JL/A connector pairs to C154 of A3, displacing its expected partner G128, which instead forms a WC pair with C111 of the JL/A connector (Fig. 3A). This base-swap is part of a broader network of tertiary interactions formed by the JL/A connector as it exits the P4 stem (Fig. 3B). A104 stacks between the terminal C103:G36 pair of P4 and A134 of the AJ3/4 loop. The C105 cytosine binds the adenine stack at A129, positioned between the ribose sugars of C130 and A135, while its 2′-hydroxyl bridges the 2′-hydroxyls of A134 and G136 within the A4 stem. This network of interactions positions the adenine stack terminus at A131 over the incoming nucleotide’s 5′-triphosphate, and a sharp turn places G132 over its nucleobase, forming the NTP binding pocket. A133 stacks between G132 and the ribose of A104, with G132 and A133 forming hydrogen bonds to the 2′-hydroxyls of C38 and C103 of the P4 stem.

Downstream of the NTP binding pocket, unpaired template nucleotides are pre-ordered by the ribozyme via the base-stacking platform within the JL/A connector. This region consists of contiguous nucleotides A108–C111, with A108 stacking onto the nucleobase at the +4 position of the template strand (Fig. 3A-B). Notably, previous crosslinking studies identified tertiary contacts between the template and the JL/A connector (*15*), which now can be rationalized by the structure (fig. S4A). Although the interaction at template position +4 is readily apparent in the consensus structure, more extensive interactions between the template and JL/A connector appear to occur more transiently. Further heterogeneity analysis of the 1.15 million particle stack (fig. S5) enabled identification of additional template nucleotides that engage with the JL/A linker and adenine stack (fig. S4B). Removal of template nucleotides downstream of the +1 position consistently reduces binding of the primer-template to the ribozyme and the rate of nucleotide addition (*15*). Because the polymerase was evolved to copy a variety of RNA templates, these base-stacking interactions may serve to stabilize the downstream template strand and prevent non-productive interactions between those nucleotides and components of the active site.

It is instructive to compare the polymerase accessory domain to the P2 region of the class I ligase. Nucleotides A133, A104, and A134 of the polymerase occupy nearly identical positions as nucleotides around the P4 stem of the ligase, while the template-bound NTP has shifted closer to P4 (Fig. 3B). The adenine stack replaces the sugar-phosphate backbone of P2, but stack nucleotides A35 and A63, which flank the P3 stem of the core, adopt identical positions in the ligase and thus appear to be remnants of the core. The P2 stem was retained in early forms of the polymerase, but was removed during later rounds of evolution (*6–9*), even as biochemical and structural studies identified a critical role for the AJ3/4 loop in NTP binding throughout the evolutionary lineage (*12, 15, 17, 18*). The portion of the P2 stem exiting P4 in the ligase becomes JL/A nucleotides 104–110 in 85h34. Acquisition of the C105 and G106 mutations in this region, together with A131 in AJ3/4, coincided with both a 100-fold increase in catalytic efficiency and improved sequence generality of the polymerase (*9*). The evolutionary history of the accessory domain is complex, with parts of the adenine stack and the NTP binding pocket emerging early, and later stages of evolution refining and stabilizing the NTP pocket through new interactions with the JL/A connector.

### A new helical element binds the primer-template duplex

The primer-template duplex docks into the catalytic core of 85h34, occupying the same position as that of the P1 stem in the class I ligase structure (*13*). The 5′-helical extension, consisting of the P8 pseudoknot and processivity tag, lies parallel to the minor groove of the primer-template helix. The base of the helix is formed by the P8 pseudoknot where it joins the P7 stem and the J1/3 single-stranded region. In the ligase, the J1/3 region contains the catalytic Mg^2+^ coordination site (equivalent to nucleotides G18, A19, and C20 in 85h34), which are positioned at the ligation junction and supported by A-minor interactions (Fig. 4, A and B). In 85h34, the apex of P7 is opened, and the adjacent P8 stem consists of five WC pairs between nucleotides G84–C88 and G11–C15, the latter located just upstream of the catalytic Mg^2+^ site (Fig. 4, C and D). The sharp 90° turn from the P7 to P8 helix is supported by a non-canonical A81:A89 pair at the junction of the two helices. The processivity tag stacks onto P8, forming an extended helical axis. At the junction of those two paired regions, nucleotides A16 and A17 exit from P8 to form A-minor interactions with the primer-template duplex.

**Fig. 4.**
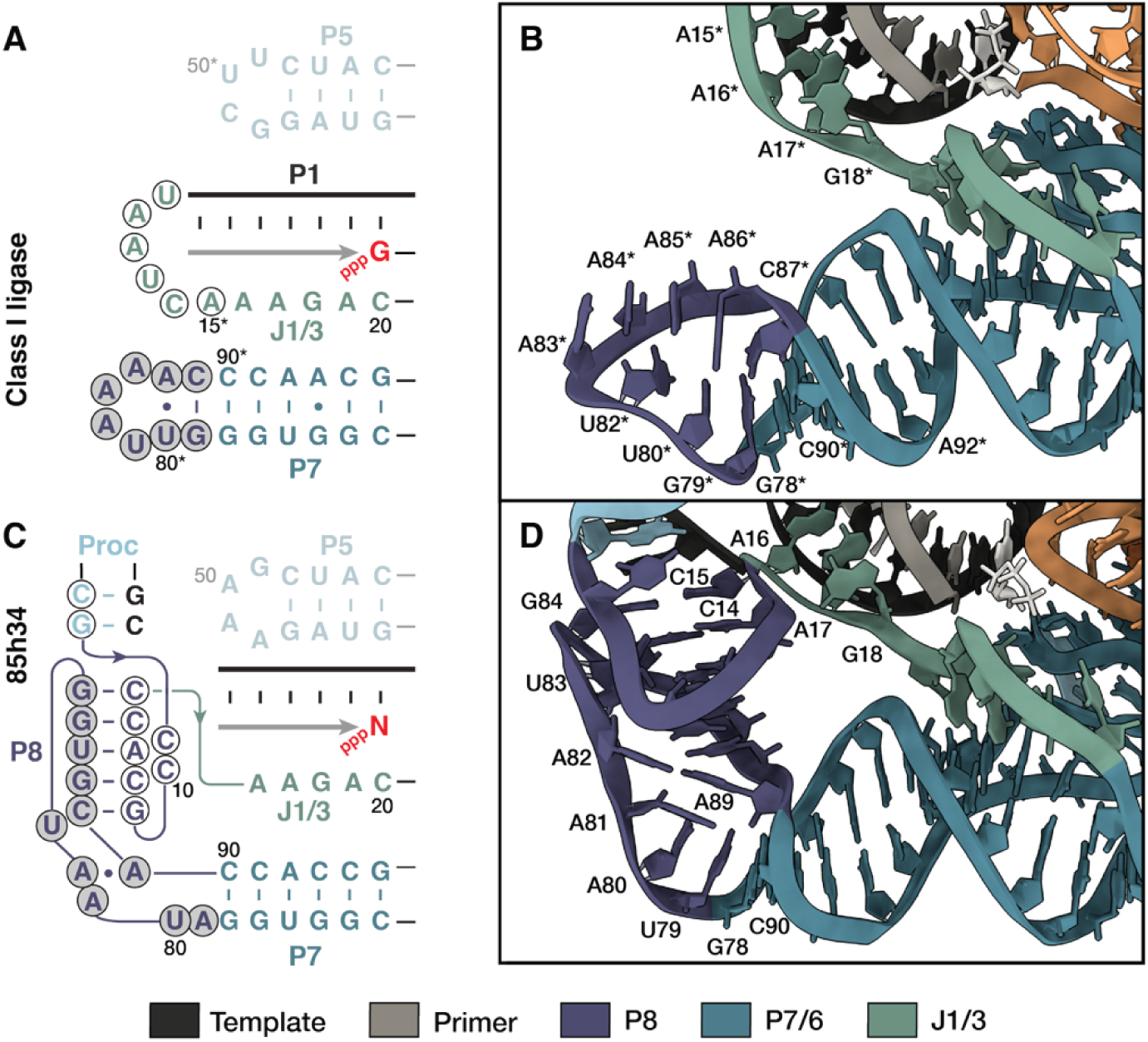
Evolution results in stabilization of the primer-template. Secondary structure diagrams showing portions of (**A**) the class I ligase and (**C**) 85h34. Nucleotides that were altered in either the J1/3 or P7 region in the ligase to form the P8 pseudoknot in 85h34 are highlighted with open or shaded circles, respectively. Atomic models focused on the J1/3, P7, and P8 regions are shown for (**B**) the class I ligase and (**D**) 85h34. Asterisks indicate adaptation of the ligase numbering convention to match that of 85h34. P8 did not exist in the class I ligase, but the region of the ligase that gave rise to P8 is highlighted.

The P8 pseudoknot was first identified in the 52-2 polymerase as a novel tertiary structural element based on mutational co-variation studies (*11*). At that time, it was noted that nucleotide A16 does not tolerate substitutions, even with a compensatory change to its potential pairing partner in U83. These results are now explained by the structure, which shows that A16 makes hydrogen-bonding contacts with the 2′-hydroxyl at position –4 of the template, a type II A-minor interaction (*19, 20*). The adjacent A17 forms a hydrogen bond with the 2′-hydroxyl at position –3 of the primer, a type III A-minor interaction. As is typical for A-minor interactions, A16 and A17 lie within a cluster supported by G18 and a Mg^2+^ ion coordinated to the phosphate of C15 (fig. S6). Deoxynucleotide substitution at either position –4 of the template or position –3 of the primer results in lower *k*_cat_ and increased *K*_m_ for nucleotide addition by the 52-2 polymerase, confirming the functional relevance of these contacts (*15*).

The P8 pseudoknot is a modular functional element that can be transplanted between polymerase variants to improve the binding constant of the ribozyme for the primer-template by more than 1000-fold (*11, 15*). By providing a docking platform for the primer-template, this domain addresses a critical shortcoming of less advanced polymerase ribozymes. Both types II and III A-minor interactions are sequence general, enabling the polymerase to bind and copy a variety of RNA templates and to operate processively along those templates. In addition, the capacity of A-minor interactions to screen for correct WC pairing within a target helix may facilitate recognition of helical distortions caused by mis-incorporated nucleotides, accounting for the observed “stalling” effect of the polymerase ribozyme when a mismatch occurs within 3– 4 nucleotides of the 3′ terminus of the product strand (*10*). Similar stalling effects have been observed in both protein polymerases (*21*) and the decoding center of the ribosome (*22*), enabling kinetic proofreading. These principles may also apply to at least one other polymerase ribozyme (*23*).

## Discussion

The evolutionary lineage leading from an ancestral polymerase that contains both the accessory domain and processivity tag to the 85h34 polymerase resulted in the acquisition of more than 40 mutations (fig. S7). Despite these changes, many structural features that are present in 85h34 were already present in that ancestor, as revealed by a recently determined *apo*-structure of tC19Z (*24*). Not only has the ligase core been preserved throughout the lineage, but also the components of the accessory domain, including the adenine stack and AJ3/4 nucleotides at the active site, occupy similar positions in both structures. The regions of structural divergence in 85h34 have, unsurprisingly, undergone the most extensive evolutionary change. These include the P8 pseudoknot and the network of tertiary contacts involving JL/A, A3, and AJ3/4 (fig. S7B). The evolutionary changes in this latter region arose early in the divergence from tC19Z and provide most of the tertiary contacts that position the adenine stack over the incoming NTP and stabilize the downstream template (fig. S7C). The changes that formed the P8 pseudoknot began at a similar point, before the 24th round of evolution, but were only fully consolidated by round 52 (*11*). The remaining changes cluster around the incoming NTP and the P3/A4 base-swap and arose later in evolution when there was strong selection pressure for improved polymerase fidelity. Mapped onto the tertiary structure of the ribozyme, evolutionary changes have primarily been directed toward positioning and stabilizing both the primer-template and incoming NTP within a catalytic core that had already been established in the class I ligase (*25*).

The structure of 85h34 provides the first glimpse into how generations of directed evolution can shape a ribozyme replication complex. Starting from the RNA ligase ancestor, distinct structural innovations helped to secure the RNA substrates within the ribozyme’s active site to enable accurate and sequence-general polymerization of RNA. These emergent architectural features are remarkably similar to those seen in single-subunit polymerase proteins, such as the RNA-dependent RNA polymerase of poliovirus (Fig. 5). When Steitz and colleagues solved the structure of the Klenow fragment of *E. coli* DNA polymerase I, they described its key features by analogy to those of a hand: the palm holds the active site, the fingers engage the incoming nucleotide, and the thumb stabilizes the extending primer-template duplex (*26*). This analogy appears to apply to the polymerase ribozyme as well: the ligase core forms the palm, the accessory domain forms the fingers that grasp the incoming NTP, and the J1/3 region, P8 pseudoknot, and processivity tag form the thumb that engages the primer-template. The overall positioning of these elements are similar among 85h34 and all four classes of single-subunit protein polymerases (fig. S8).

**Fig. 5.**
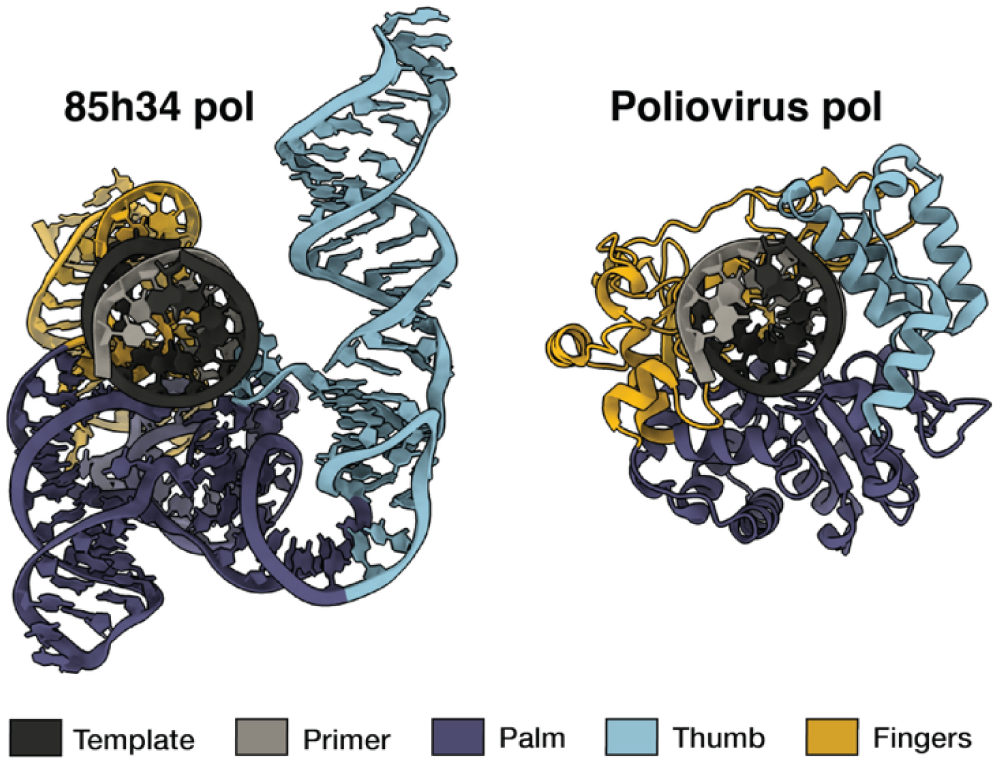
Structural convergence of the RNA-dependent RNA polymerase ribozyme and protein polymerases. The 85h34 replication complex is displayed alongside poliovirus RNA-dependent RNA polymerase protein (PDB ID 3OL6) (*32*), using the right-hand analogy. Both complexes are aligned to the active site and primer-template, as viewed down the axis of the primer-template helix.

In 1968, long before the discovery of catalytic RNA, Francis Crick suggested that “possibly the first ‘enzyme’ was an RNA molecule with RNA replicase properties” (*1*). This notion became more tangible following the discovery of naturally-occurring ribozymes that catalyze nucleotidyl transfer reactions (*27*) and the directed evolution of RNA ligase and RNA polymerase ribozymes (*3, 6*). Crick also said that “tRNA looks like Nature’s attempt to make RNA do the job of a protein” (*28*). With regard to both structure and mechanism, the 85h34 polymerase demonstrates evolution’s ability to make RNA do the job that modern life has assigned to proteins. Because the ribozyme was evolved to copy RNA, with no design or selective step favoring the hand architecture, its emergence may reflect a general model for copying the double helix. It appears that the blind watchmaker of evolution, through necessity and not foresight, can find a common path home.

## Materials and Methods

### Preparation of RNA

The sequences of all oligonucleotides used in this study are listed in table S2. RNA primers P1 and P1fb were prepared by solid-phase synthesis using an Expedite 8909 DNA/RNA synthesizer, with reagents and phosphoramidites from either Glen Research or ChemGenes. Template T1 and 85h34 polymerase RNA were prepared by *in vitro* transcription in a mixture containing 5–20 ng/μL template DNA, 5 mM each NTP, 15 U/μL T7 RNA polymerase, 0.002 U/μL inorganic pyrophosphatase, 25 mM MgCl_2_, 2 mM spermidine, 10 mM DTT, and 40 mM Tris-HCl (pH 8.0), which was incubated at 37 °C for 2 h. The template DNA was then digested by adding 0.1 U/μL Turbo DNase (Thermo Fisher) and incubating at 37 °C for 1 h. Synthetic oligodeoxynucleotides were obtained from Integrated DNA Technologies (IDT). For preparation of template 1, the corresponding oligodeoxynucleotides were annealed and used directly. For preparation of the 85h34 polymerase, the corresponding portion a plasmid encoding the 85h34 sequence was PCR amplified using primers Fwd1 and Rev1 and Q5 Hot Start polymerase (New England Biolabs). All RNAs were purified by denaturing polyacrylamide gel electrophoresis (PAGE) prior to use. NTPs and α,β-methylene ATP (AMPcPP) were from Jena Biosciences, and all other reagents were from Sigma-Aldrich.

### Assay of polymerase activity

Primer extension reactions were conducted in a mixture containing 1 µM 85h34 polymerase, 1 µM primer P1fb, 1 µM template T1, 2 mM each NTP, 50 mM MgCl_2_, 50 mM Tris-HCl (pH 8.3), and 0.05% (w/v) TWEEN20, which was incubated at 25 °C. Prior to initiating the reaction, the RNA components were folded by incubating at 80 °C for 30 s, slowly cooling to 25 °C, adding 50 mM MgCl_2_, and incubating at 25 °C for 10 min. The reaction was initiated by adding the NTPs and quenched by adding Na_2_EDTA. The extended products were purified by biotin capture on MyOne C1 streptavidin magnetic beads (Thermo Fisher) as previously described (*12*), then separated by PAGE and imaged using an Amersham Typhoon RGB laser scanner and quantified using ImageQuantTL software.

### Assembly of polymerase ribozyme replication complex

The replication complex was assembled by mixing 1 μM each of 85h34 polymerase, primer P1, and template T1 in the presence of 10 mM Tris-HCl (pH 8.3) and 1 mM Na_2_EDTA, which were heated at 80° C for 30 s, then cooled at 0.2 °C/s to 25° C. 5 mM AMPcPP and 50 mM MgCl_2_ then were added to the mixture to complete the holoenzyme. Assembled complexes were held at 25 °C for 10 min, then diluted with the same solution to 10 nM concentration of each RNA, and immediately transferred to MassGlass NA coverslips for data collection on the Refeyn 2MP Mass Photometery instrument. Movies (60 s) were acquired with AcquireMP and analyzed with DiscoverMP software (Refeyn) using default settings to obtain distribution histograms of event counts and Gaussian fits to individual peaks. Complexes for cryo-EM analysis were prepared in the same manner, but using 2 μM each of 85h34, P1, and T1, 75 mM MgCl_2_, and 7.5 mM AMPcPP. After incubating the mixture at 25 °C for 10 min, 500 μL of solution was concentrated to 100 µL using a Pierce Protein Concentrator PES, 30K MWCO. A portion of the complex was diluted and assayed by mass photometry, with the remainder used for cryo-EM grid preparation.

### Cryo-EM specimen preparation

Samples were prepared by spiking 1 μL of *n*-decyl-β-D-maltopyranoside (DM) detergent (*33*) into 9 μL of 10 μM replication complex. Final detergent concentrations ranged from 0.05% to 0.002% DM. UltrAufoil R0.6/1.0 grids (Quantifoil) were hydrophilized via plasma comprised of 35.0 sccm argon and 11.5 sccm oxygen at 40 W for 40 s using a Solarus II Model 955 plasma cleaning instrument (Gatan). Grids were immediately loaded onto a Vitrobot Mark IV (Thermo Fisher), equilibrated at 25 °C with 100% humidity. 3.5 μL of sample were loaded onto the grid, blotted for 6 s with blot force set to 1, and immediately plunge froze into liquid ethane. Grids were clipped into autogrid cassettes under liquid nitrogen.

### Data collection

Screening of grids and automated data collection were performed on a FEI Titan Krios G3 instrument (Thermo Fisher) equipped with a K3 direct electron detector (Gatan). Several grids prepared from samples containing DM concentrations of 0.0017% or 0.006% were identified as suitable for data collection based on the presence and distribution of appropriately sized particles. Automated data collection of 50-frame movies was carried out using a total dose of 50 electrons/Å^2^ over a 2.7 s exposure with a pixel size of 0.83 Å, resulting in a dose rate of 0.69 electrons/pixel/frame. Defocus was randomly applied between the range of –0.7 to –2.5 μm during collection. The cryo-EM dataset consisted of 34,997 movies (table S1).

### Data processing

An overview of the data processing scheme is shown in fig. S2. CryoSPARC version 4.7.0 (Structura Biotechnology) was used for pre-processing, particle picking with templates and with topaz (*34*), *ab initio* reconstructions, heterogeneous 3D classification, initial high-resolution refinement, and final map quality control analyses. Relion version 5.0 was used for the final high-resolution refinement and focused 3D classification with blush regularization. A consensus map was generated using a particle stack of 1.15 million particles. 3D classification with alignment was performed using heterogeneous refinement in CryoSPARC, but this strategy was not used for final reconstructions. Instead, an active site focus mask was used to identify 5 subsets of particles with distinct conformational states. Important parameters for 3D classification were disabling 3D alignment, increasing the regularization parameter T to 50, and enabling blush regularization (*35*). The final particle stacks were imported into CryoSPARC for reconstruction, gold standard global Fourier shell correlation (FSC) analysis, 3D FSC analysis (*36*), and sampling compensation factor calculation (*37*, *38*). The final consensus map was resolved to 2.8 Å. The resolution assessment is consistent with map features that show separation of nucleobases within helical stacks, clearly distinguishable densities for sugars, phosphates, and nucleobases, and ordered waters and Mg^2+^ ions (fig. S2B). All figures for the main text were generated using models derived from the consensus map.

Following derivation of the consensus map and atomic model, efforts were made to identify conformational changes and alternative active site configurations. Heterogeneity analyses using 3D classification with alignment in CryoSPARC separated particle stacks based on dynamics corresponding primarily to the termini of the processivity tag and primer-template duplex and to the 3’ end of the polymerase, which are known to be non-essential for function (fig. S5, A and B). Regions that contributed to global heterogeneity were masked out and focused classification was used to sort the particles into 10 classes and reconstructed maps of the 85h34 core to identify changes in the active site (fig. S5, C and D). However, modeling into the resulting maps did not reveal significant conformational changes compared to the consensus map for the 85h34 polymerase. Rather, structural changes were noted in the template RNA near the JL/A loop and the top of the adenine stack. In two of the subclasses (subclasses 2 and 7), it was possible to model the density as distinct dinucleotides within the template, consistent with published cross-linking data (fig. S4; fig. S5 E and F), and these maps and models are accordingly deposited alongside the consensus structures (table S1).

### Model building and refinement

An initial model of the replication complex was predicted using Alphafold3 (*39*). In general, the predictions matched poorly to the experimental density, with the exception that the catalytic core resembled the structure of the class I ligase (*13*) and could easily be docked into the experimental consensus map. The rest of the replication complex model was built manually by tracing the backbone phosphates leading away from the catalytic core using WinCoot v1.3.1 (*40*). Mg^2+^ ions were identified by eye and modeled manually based on the local chemical environment. Waters were automatically modeled using *phenix.douse* with *keep_input_water*=true, *max_dist*=1.9, *sphericity_filter*=false, and *map_threshold* set to 5 RMSD of each map. Waters were inspected manually. Metal coordination restraints for real space refinement were generated using *phenix.ready_set* with *output_angles*=true. The final model was then refined in real space using Phenix v1.21 (*41*). The model was inspected by hand and with the aid of MolProbity (*42*). The model was also used as a starting point to refine models corresponding to subclasses 2 and 7, which was performed using similar procedures. Images of maps and models were rendered using Chimera v1.17.3 (*43*) and ChimeraX v1.12 (*44*).

## Supplementary Text

There is insufficient structural information to determine whether the transition state for phosphodiester bond formation proceeds through the classical two-metal ion mechanism that is ubiquitous for protein polymerases. This mechanism entails a trigonal planar configuration of the α-phosphate with two coordinated Mg^2+^ ions (termed metal A and metal B) that are spaced ∼4 Å apart. In the 85h34 structure, the Mg^2+^ that coordinates the A19 phosphate is presumed to be metal A. There is a nearby Mg^2+^ coordinating the β,γ-phosphates of the incoming NTP to the phosphate of A21, but that Mg^2+^ is 7 Å away from metal A. Bartel and colleagues proposed that N4 of C47 in the ligase (C37 in the polymerase) serves in place of metal B, coordinating the α-phosphate of the incoming NTP to achieve a similar transition state geometry (*13*). Kinetic studies of an earlier form of the polymerase demonstrated a linear free energy relationship for the ribozyme polymerase similar to that of protein polymerases (*45*). The 85h34 structure cannot distinguish between the two transition state models. It is possible that the choice of non-reactive NTP analog caused the active site to be in a non-productive state. Structures of protein polymerases bound with α,β-methylene NTP analogs have been found to trap the active site in an “open”, non-catalytic configuration (*46*, *47*).

The 85h34 structure is also unable to explain the mechanisms by which the polymerase discriminates between WC paired and mismatched NTPs within the active site. The nucleotide at position 132 does appear to gate selectivity in the NTP binding pocket. The previously conserved A132 has mutated to G in the most advanced forms of the polymerase that were under strong selection pressure for improved fidelity, including 85h34 (*12*). Mutation of A132 to U in another polymerase variant was associated with relaxed substrate selectivity (*18*). However, further studies are needed to determine the mechanisms by which G132 and other nucleotides forming the NTP binding pocket might screen for WC pairing between the NTP and corresponding template nucleotide.

## Acknowledgments

We thank Deni Szokoli, Jason Hingley, Navtej Toor, Rhiju Das, Jack Szostak, and Joseph Picirilli for providing helpful feedback on our preliminary structural analyses. A portion of this research was supported by NIH grant R24GM154185 and performed at the Pacific Northwest Center for Cryo-EM (PNCC) with assistance from Sean K. Mulligan.

## Author contributions

Conceptualization: TSS, WGC, DPH, DL, GFJ

Methodology: TSS, DPH, WGC, XH, DL

Investigation: TSS, DPH, LA, XH, WGC

Visualization: TSS, DPH

Funding acquisition: TSS, DPH, WGC, GFJ, DL

Project administration: GFJ, DL

Supervision: GFJ, DL

Writing – original draft: TSS

Writing – review & editing: TSS, DPH, DL, GFJ

## Data, code, and materials availability

The resulting coulombic potential maps derived from our cryogenic electron microscopy analyses are available at EMDB entries EMD-78851, EMD-78852, and EMD-78853. Atomic models are available at PDB entries pdb_000038JC, pdb_000038JD, and pdb_000038JE. Other data, code, and analysis will be made available upon request within reason.

---

**Fig. S1.**
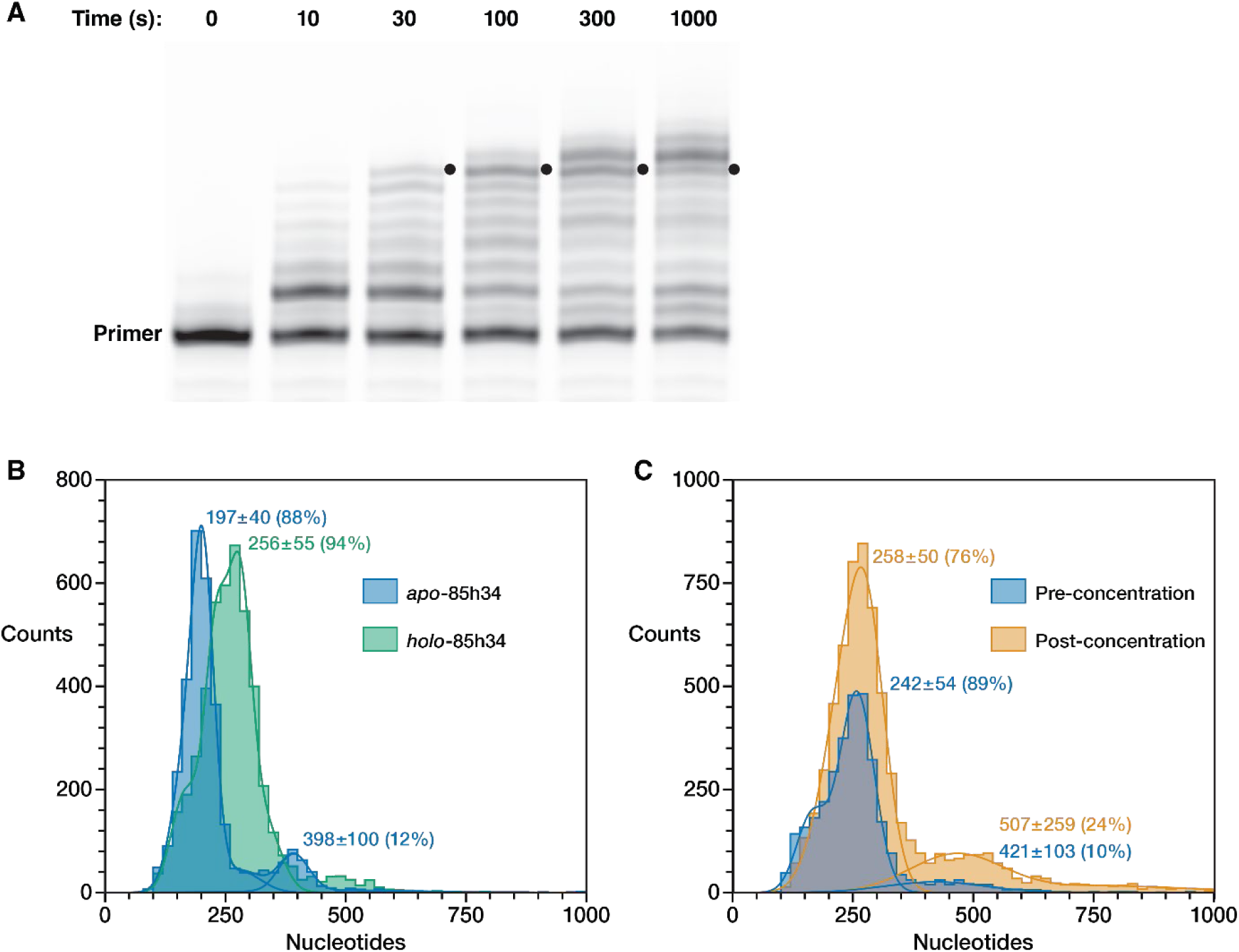
Biochemical characterization of 85h34. (**A**) Activity of the 85h34 polymerase, extending 5’-fluorescein-labeled primer P1 on template T1 in the presence of 2 mM each NTP and 50 mM MgCl_2_ at pH 8.3 and 25 °C. The extension products were separated by PAGE. Black dots indicate full-length products. (**B**) Replication complexes were assembled using 85h34, P1, T1 and AMPcPP and analyzed by mass photometry. The complex peak is compared to 85h34 alone. (**C**) The assembled complexes were concentrated 10-fold using a spin-concentration device, then re-analyzed by mass photometry prior to vitrification for cryo-EM analysis. The expected sizes of monomeric *apo*- and *holo*-85h34 are 194 and 257 nucleotides, respectively. Individual peaks are labeled with the calculated mean and standard deviation of the nucleotide content based on a Gaussian fit of the data, together with the fraction of the population in that peak.

**Fig. S2.**
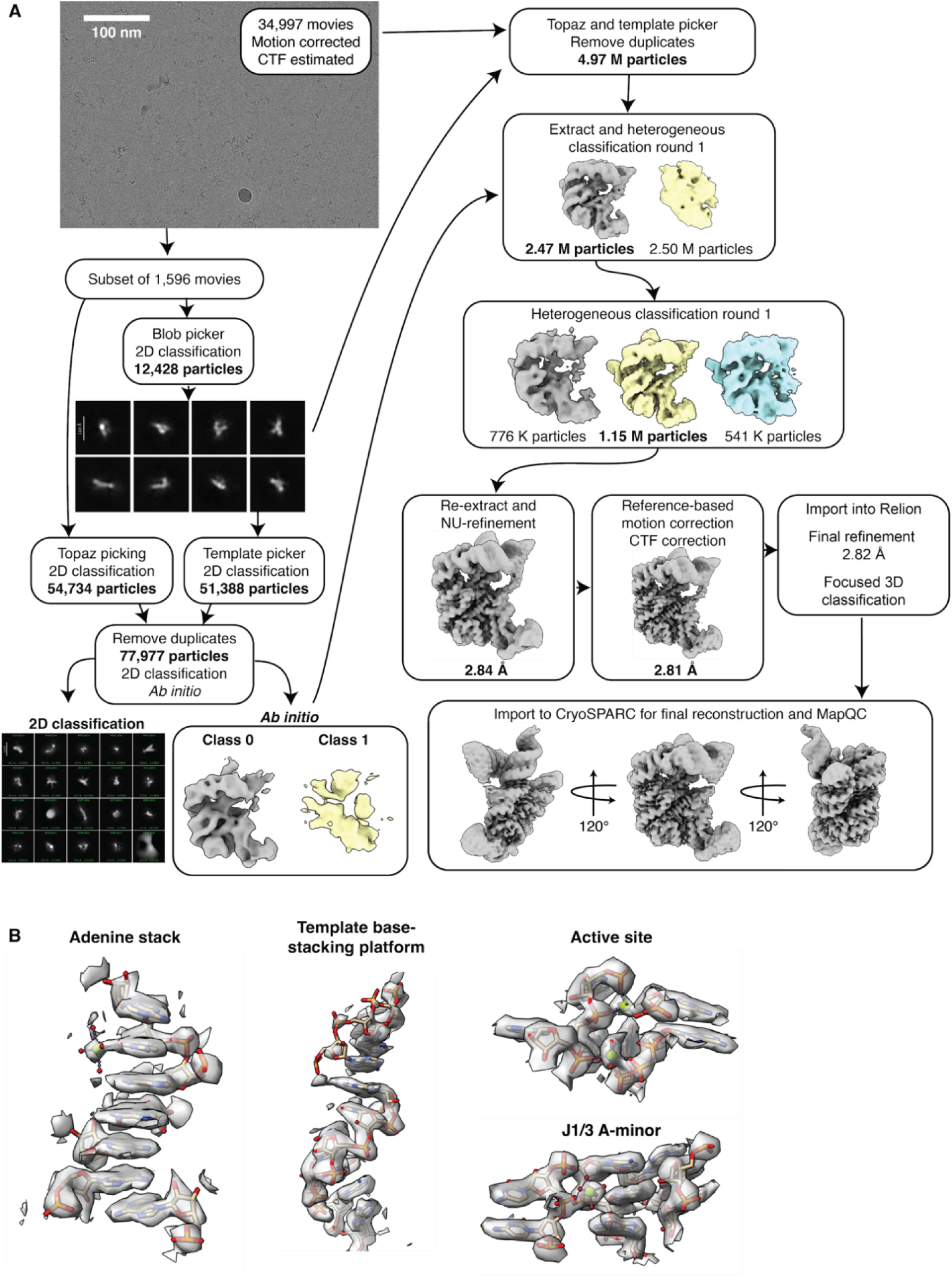
Cryo-EM data processing scheme. (**A**) Representative micrographs, 2D classification results, and representative coulombic potential maps. (**B**) Snapshots of models fitted within the sharpened map.

**Fig. S3.**
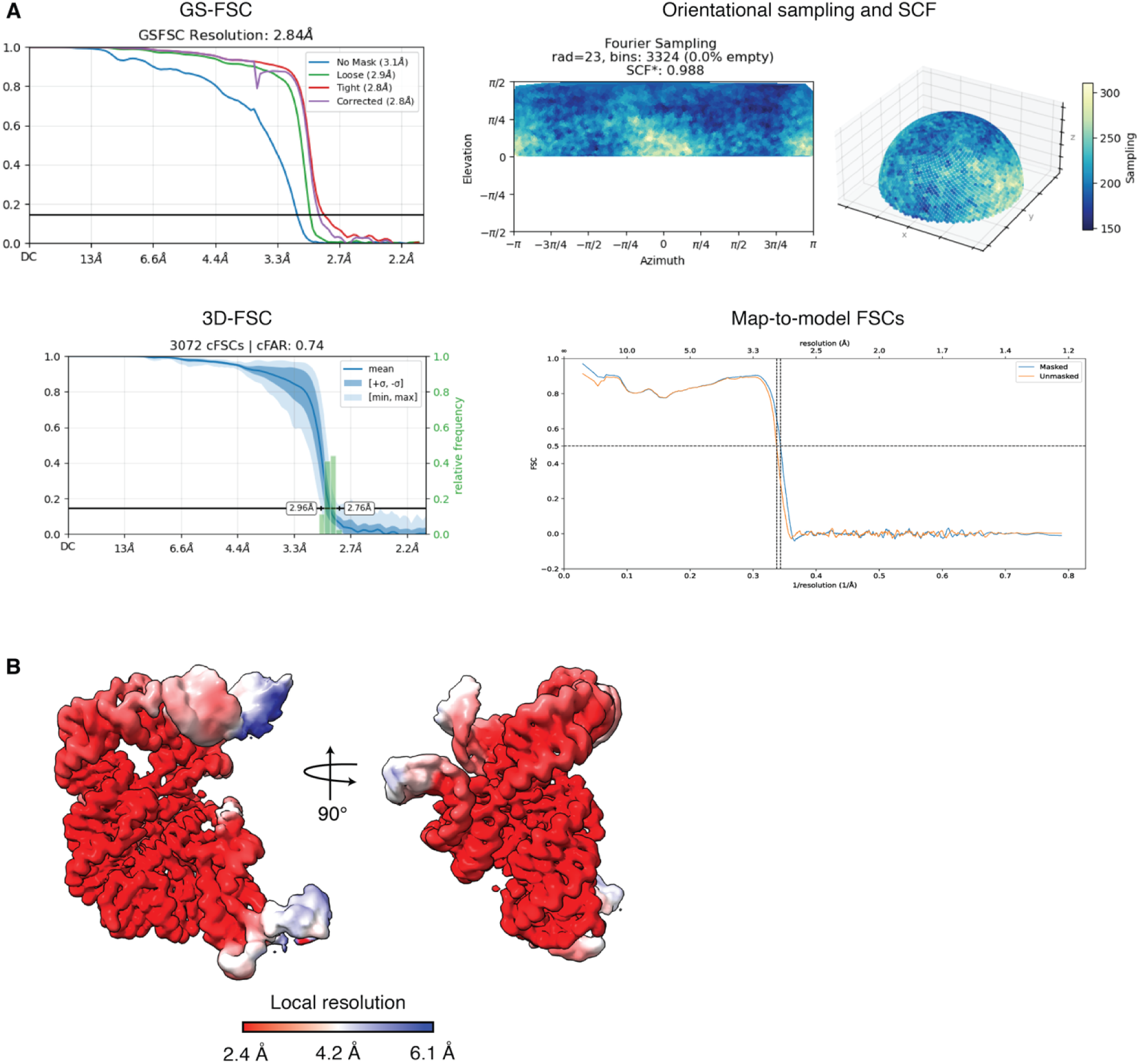
Cryo-EM map quality control measurements. (**A**) Grid showing gold standard Fourier shell correlations (GS-FSC), 3D conical FSCs (3D-FSC), orientational sampling as depicted as an Euler plot and on a half globe, sampling compensation factor and empty Fourier bins calculations, and map-to-model FSCs. (**B**) Local resolution was painted onto the coulombic potential map according to the color key.

**Fig. S4.**
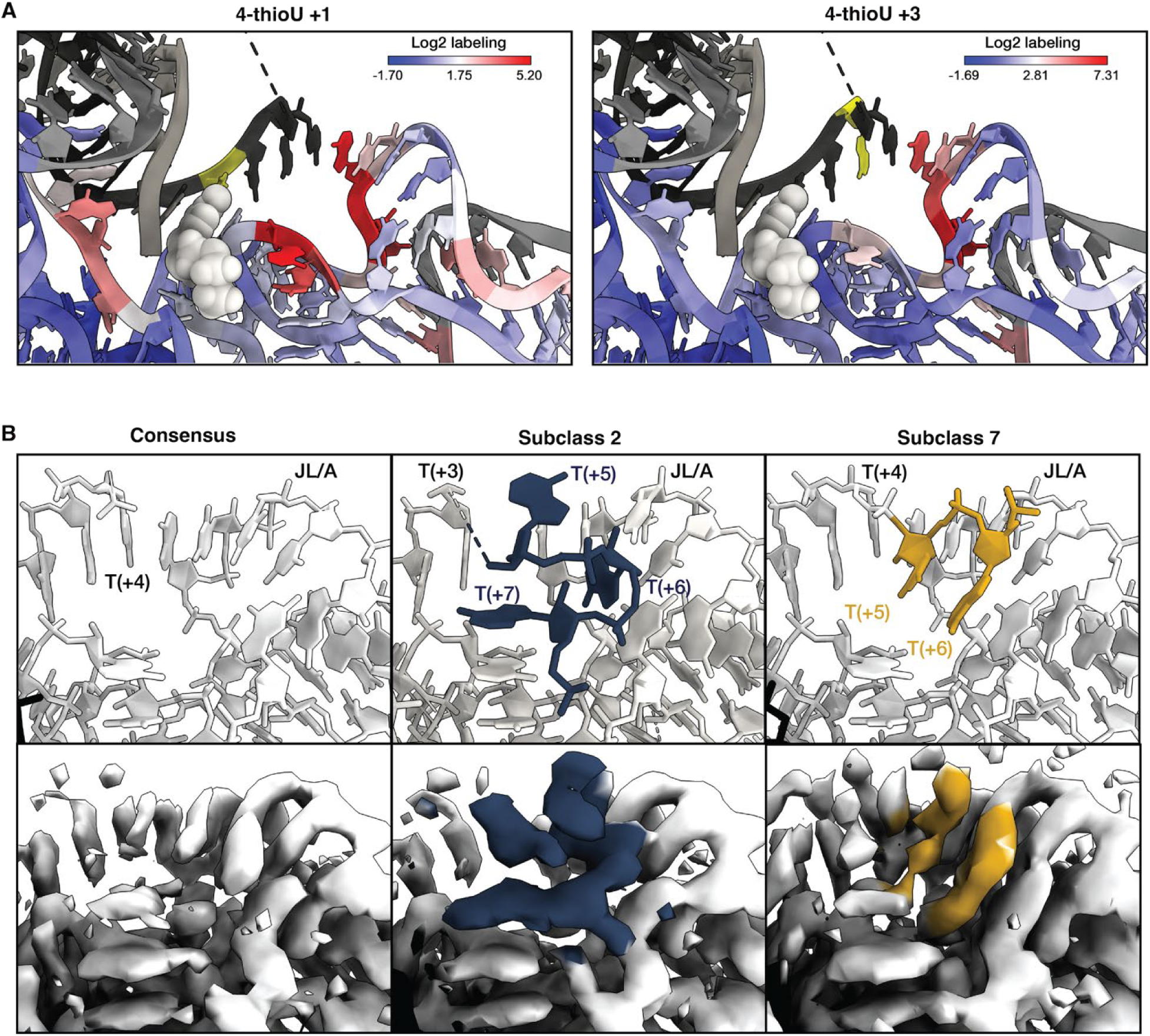
Tertiary contacts between the template and accessory domain. (**A**) Labeling of cross-linked positions in the ribozyme based on a previous study in which 4-thioU was installed at either the +1 or +3 position of the template and UV-induced cross-links were identified by stalling of reverse transcriptase (*15*). Cross-linking efficiency at various nucleotide positions is shown on a log_2_ scale. Positions without values are shown in gray. The position where 4-thioU was installed is colored yellow. AMPcPP is shown as space filling atoms, colored white. (**B**) Structural detail at the site of cross-linking, comparing the consensus map and two other subclasses that were identified by focused 3D classification (fig. S5). Significant heterogeneity is seen in the downstream portion of the template (positions +4 to +7) and the base-stacking platform within the JL/A loop.

**Fig. S5.**
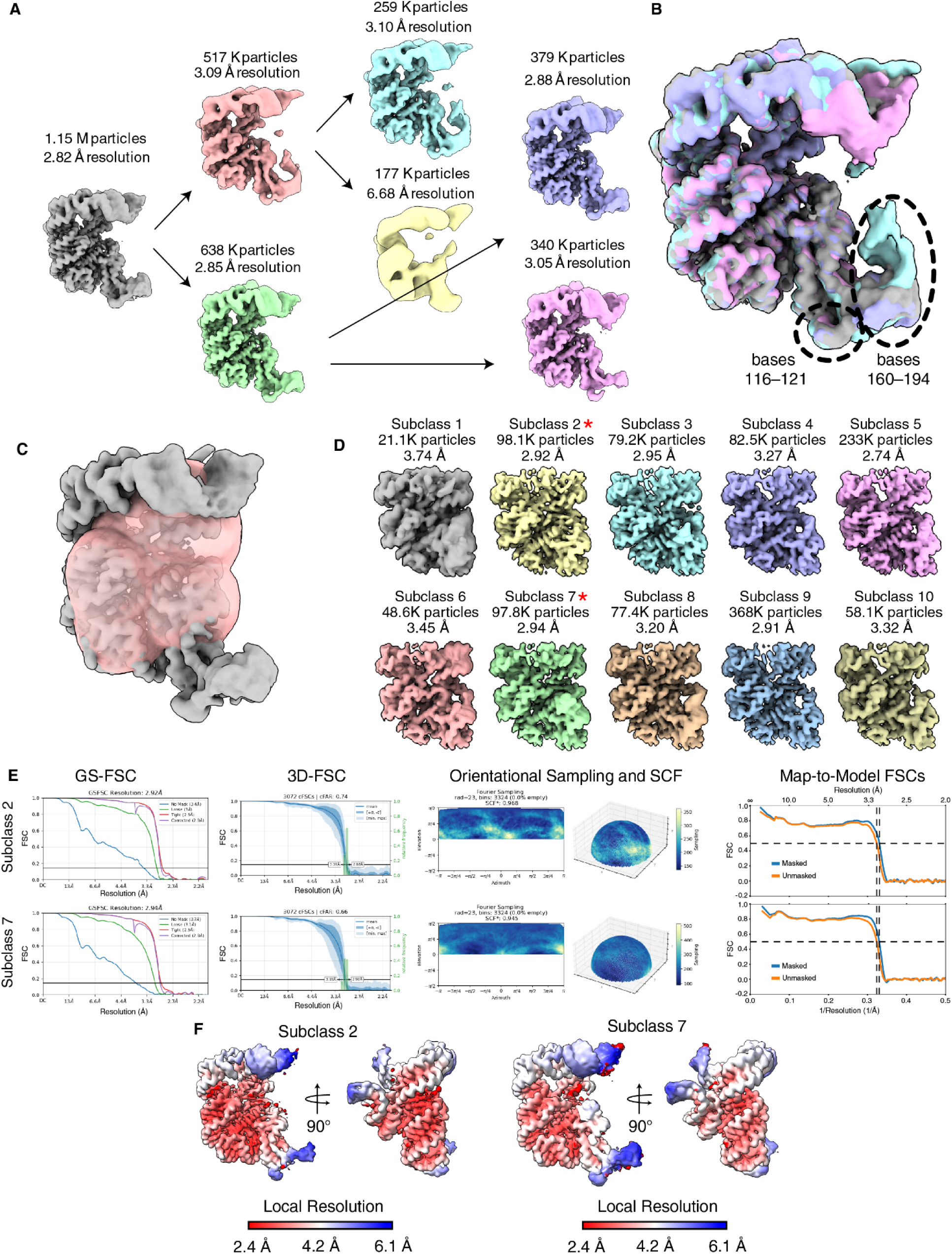
Heterogeneity analysis of consensus particle stack. (**A**) Global heterogeneity was assessed using iterative 3D classification with alignment into two classes. (**B**) Resulting maps are overlayed with the consensus map. (**C**) Masking strategy focusing on the active site for focused 3D classification without alignment. (**D**) 3D classification results for 10 classes are shown. The resulting maps did not reveal any notable differences. However, subclasses 2 and 7 (red star) contain additional density corresponding to downstream portions of the template strand, which could reliably be interpreted with an atomic model and is supported by previous cross-linking studies (fig. S4) (*15*). Although subclass 2 was nominally resolved to 2.74 Å, which is higher than the consensus map, it did not reveal any additional features within the active site. Because the consensus map showed more evenly distributed resolution, it was used for all analyses in this study. (**E**) Grid showing gold standard Fourier shell correlations (GS-FSC), 3D conical FSCs (3D-FSC), orientational sampling as depicted as an Euler plot and on a half globe, sampling compensation factor and empty Fourier bins calculations, and map-to-model FSCs. (**F**) Local resolution was painted onto the coulombic potential map according to the color key.

**Fig. S6.**
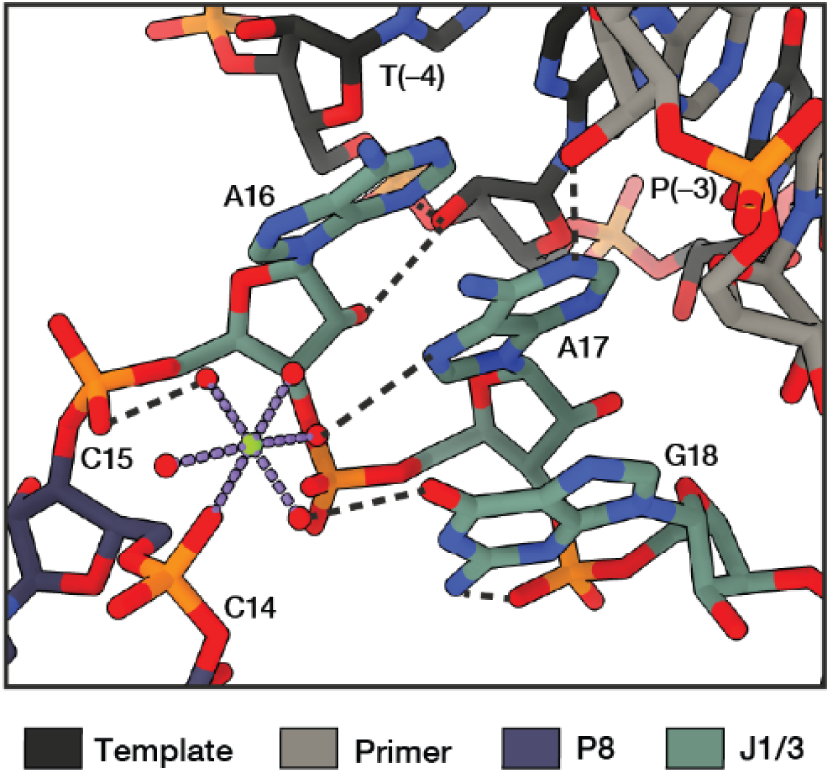
J1/3 A-minor tertiary contacts with the primer-template. Atomic model of J1/3 A-minor interactions, with hydrogen bonds and metal coordination shown as dashed black and purple lines, respectively.

**Fig. S7.**
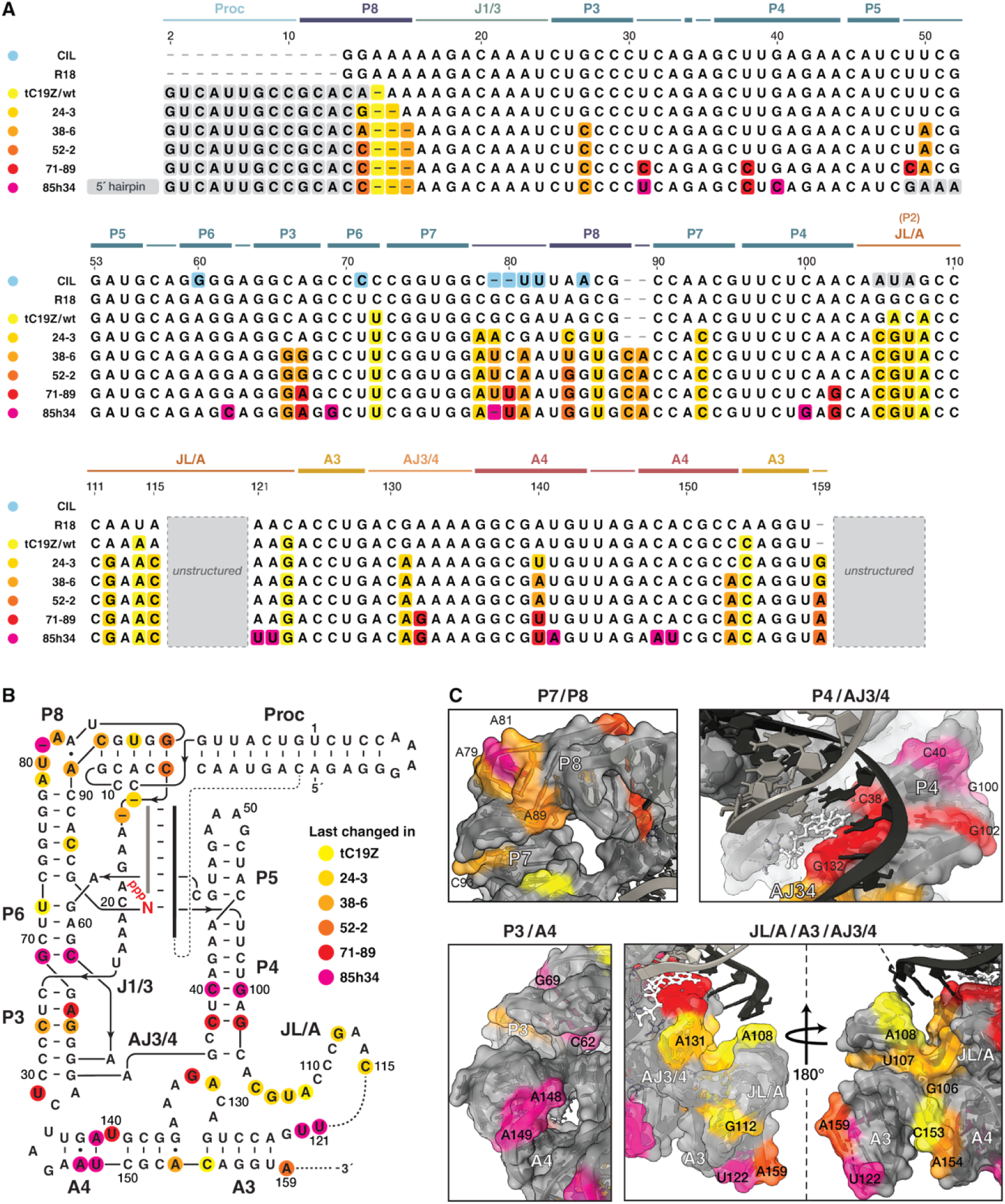
Structural analysis of the mutational landscape. (**A**) Sequence alignment of 85h34 and ancestral polymerase ribozymes: the class I ligase (CIL) (*5*, *6*); the first polymerase ribozyme, R18 (*6*); the first polymerase to contain the processivity tag, tC19Z (*8*); and polymerases from rounds 24, 38, 52, 71 in the evolutionary lineage from tC19Z to 85h34 (*9–12*), The tC19Z sequence includes mutations introduced at the outset of the 85h34 lineage (*9*). Mutations are color coded based on the generation of evolution in which they first appeared. (**B**) Mutations are highlighted in the secondary structure diagram and (**C**) painted onto snapshots of the structure of the 85h34 replication complex. Where multiple mutations occurred at the same site, mutations are colored by the last change that occurred at the position.

**Fig. S8.**
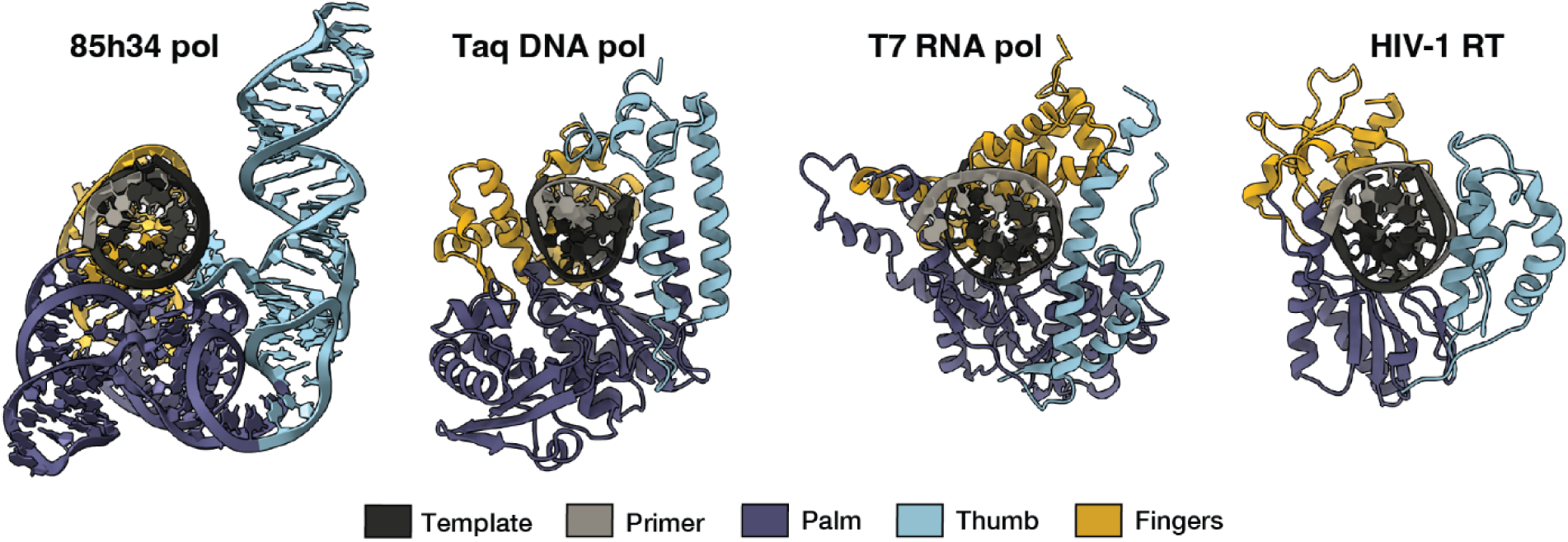
Structural convergence of the ribozyme replication complex with protein polymerases. The 85h34 replication complex structure is contrasted with protein holoenzyme complexes, including Taq DNA polymerase PDBID 1TAU (*48*), T7 RNA polymerase PDBID 1S76 (*30*), and HIV-1 reverse transcriptase PDBID 2HMI (*49*).

**Table S1.** Cryo-EM data collection, processing, and model building statistics.

| Data collection |  | Holo-85h34 replication complex |  |  |  |  |  |
| --- | --- | --- | --- | --- | --- | --- | --- |
| Microscope |  | Titan Krios G3 (Thermo Fisher) |  |  |  |  |  |
| Detector |  | K3 (Gatan) |  |  |  |  |  |
| Voltage (source) |  | 300 kEV |  |  |  |  |  |
| Magnification |  | 105000X |  |  |  |  |  |
| Pixel size |  | 0.83 Å |  |  |  |  |  |
| Total dose |  | 50.0 e-/Å2 |  |  |  |  |  |
| Exposure time |  | 2.7 seconds |  |  |  |  |  |
| Frames |  | 50 |  |  |  |  |  |
| Dose rate |  | 0.69 e-/pixel/frame |  |  |  |  |  |
| Number of frames |  | 50 |  |  |  |  |  |
| Defocus range |  | −0.7 to −2.5 μm |  |  |  |  |  |
| Automation software |  | EPU |  |  |  |  |  |
| Movies collected |  | 34997 |  |  |  |  |  |
| Data Processing |  | Consensus map |  | Subclass 2 |  | Subclass 7 |  |
| Data processing software |  | CryoSPARC v4.7.0 /<br>Relion v5.0b |  | CryoSPARC v4.7.0 /<br>Relion v5.0b |  | CryoSPARC v4.7.0 /<br>Relion v5.0b |  |
| Total extracted particles |  | 4854135 |  | 1155053 |  | 1155053 |  |
| Final particle stack |  | 1155053 |  | 98137 |  | 97830 |  |
| Gold standard FSC |  | 2.84 Å |  | 2.92 Å |  | 2.94 Å |  |
| 3D FSC range |  | 2.92–2.45 Å |  | 3.07–2.88 Å |  | 3.19–2.90 Å |  |
| Sampling compensation factor |  | 0.99 |  | 0.97 |  | 0.95 |  |
| EMDB ascension |  | EMD-78851 |  | EMD-78852 |  | EMD-78853 |  |
| Modeling |  |  |  |  |  |  |  |
| Software |  | Coot 1.2 / Phenix 2.1 |  | Coot 1.2 / Phenix 2.1 |  | Coot 1.2 / Phenix 2.1 |  |
| Identifier |  | pdb_000038JC |  | pdb_000038JD |  | pdb_000038JE |  |
| Map-to-model FSC (masked, cut off 0.5) |  | 2.93 Å |  | 3.03 Å |  | 3.03 Å |  |
| All-atom | Clashscore, all atoms: | 2.89 |  | 1.67 |  | 3.64 |  |
| Contacts | Wrong sugar puckers: | 0 | 0.5% | 0 | 0.0% | 3 | 1.8% |
| Nucleic acid | Bad backbone: | 39 | 16.5% | 35 | 16.3% | 69 | 33.6% |
| Geometry | Bad bonds: | 0 / 5128 | 0.0% | 0 / 5175 | 0.0% | 0 / 4891 | 0.0% |
|  | Bad angles: | 0 / 7979 | 0.0% | 0 / 8052 | 0.0% | 0 / 7607 | 0.0% |
|  | Chiral volume outliers: | 0/1064 | 0.0% | 0/1074 | 0.0% | 0/1014 | 0.0% |
| Composition | Nucleic acid | 213 |  | 221 |  | 203 |  |
|  | Waters | 198 |  | 155 |  | 82 |  |
|  | Magnesium | 14 |  | 13 |  | 13 |  |
|  | Waters with clashes | 9 | 4.5% | 12 | 7.7% | 1 | 1.2% |

**Table S2.**
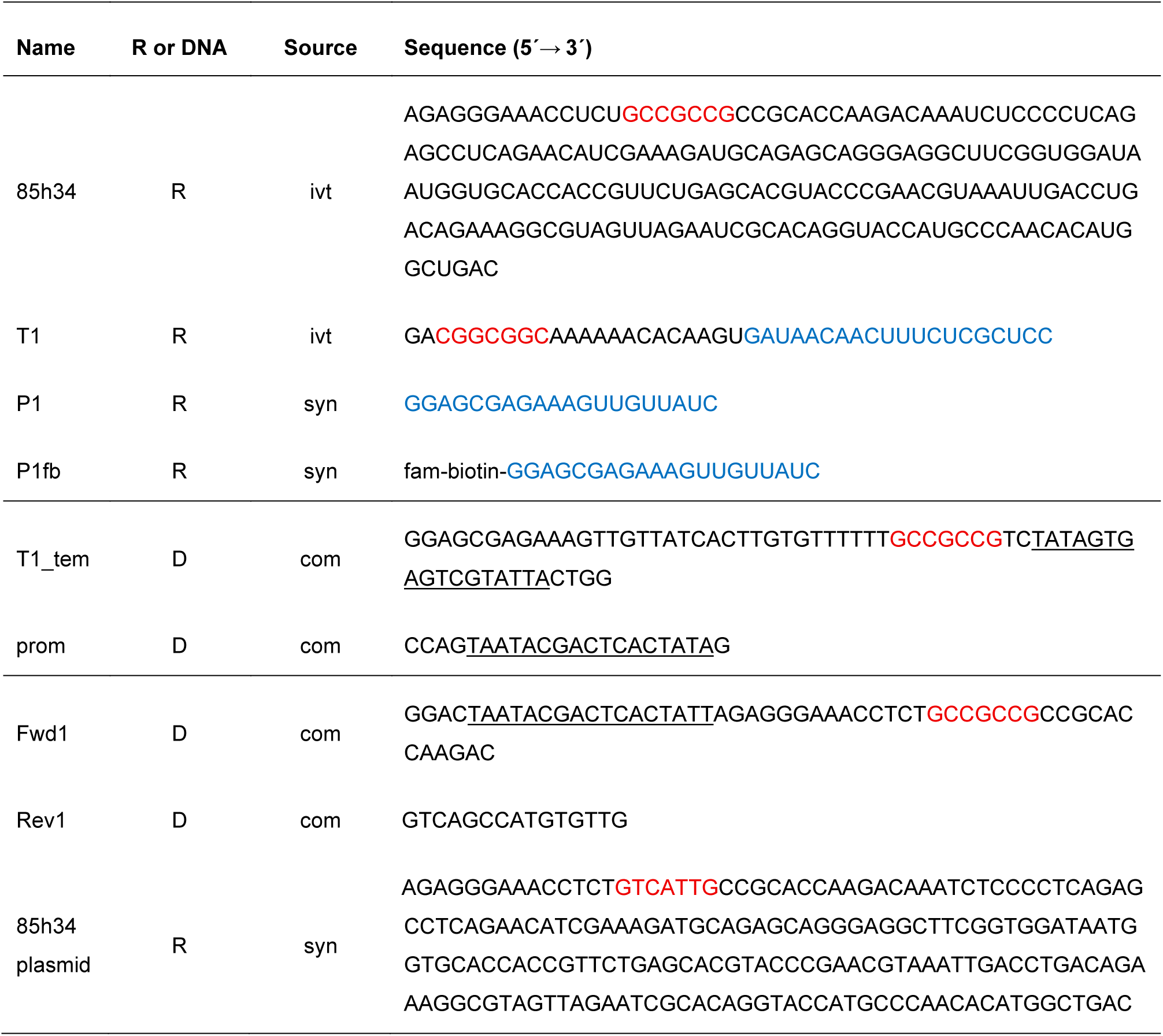
Sequences of RNA and DNA molecules used in this study. Oligonucleotides were either synthesized in-house (syn), purchased from IDT (com), or prepared by *in vitro* transcription (ivt). The 5′ end of P1fb was modified with both fluorescein (fam) and biotin. DNA encoding the 85h34 sequence was purchased from IDT as a “Rapid Gene” pUCIDT (Kan) plasmid. Sequences in red indicate the processivity tag within the ribozyme and RNA template. Note this sequence differs in materials used for cryo-EM analysis compared to the plasmid. Complementary regions between the primer and template are shown in blue. The T7 RNA polymerase promoter sequence within the forward primer or template is underlined.

